# Regions of visual cortex responding to tactile stimulation in an individual with longstanding low vision are not causally involved in tactile processing performance

**DOI:** 10.1101/2021.05.18.444648

**Authors:** Edward H. Silson, Andre D. Gouws, Gordon E. Legge, Antony B. Morland

## Abstract

Braille reading and other tactile discrimination tasks recruit the visual cortex of both blind and normally sighted individuals undergoing short-term visual deprivation. Prior functional magnetic resonance imaging (fMRI) work in patient ‘S’, a visually impaired adult with the rare ability to read both highly magnified print visually and Braille by touch, found that foveal representations of S’s visual cortex were recruited during tactile perception, whereas peripheral regions were recruited during visual perception. Here, we test the causal nature of tactile responses in the visual cortex of S by combining tactile and visual psychophysics with repetitive transcranial magnetic stimulation (rTMS). First, we replicate this prior fMRI work in S. Second, we demonstrate that transient disruption of S’s foveal visual cortex has no measurable impact on S’s tactile processing performance compared to that of healthy controls – a pattern not predicted by the fMRI results. Third, stimulation of foveal visual cortex maximally disrupted visual processing performance in both S and controls, suggesting the possibility of preserved visual function within S’s foveal cortex. Finally, stimulation of somatosensory cortex induced the expected disruption to tactile processing performance in both S and controls. These data suggest that tactile responses in S’s foveal representation reflect unmasking of latent connections between visual and somatosensory cortices and not behaviourally relevant cross-modal plasticity. Unlike studies in congenitally blind individuals, it is possible that the absence of complete visual loss in S has limited the degree of causally impactful cross-modal reorganisation.

**Significance statement:** Prior fMRI work in patient ‘S’ identified that foveal portions of S’s visual cortex respond more to tactile processing, whereas peripheral portions respond more to visual processing. Here, we tested whether this foveal processing was causally related to either tactile or visual processing. First, using fMRI we replicate prior work. Second, we demonstrate that TMS of the foveal representation and of somatosensory cortex interfered with visual and tactile discriminations respectively in controls and crucially also in S. The foveal representation in S, which is responsive to tactile stimulation, does not however play a causal role in mediating S’s ability to discriminate Braille characters and likely reflects the unmasking of latent connections between visual and somatosensory cortices.

## Introduction

Whether or not human visual cortex reorganises functionally following deprived visual input is a crucial question in visual neuroscience (Baseler et al., 2011; Cheung et al., 2009; Haak et al., 2015; Sadato et al., 1996). In blind individuals, fMRI studies highlight visual cortex activity during somatosensory tasks (Burton et al., 2002; Sadato et al., 1996) (e.g. Braille reading) and short-term visual deprivation can lead to increased recruitment of visual cortex during somatosensory tasks in normally sighted individuals (Kauffman et al., 2002; Merabet et al., 2007, 2008). Further, somatosensory and auditory task-related activity has been reported in the lesion-projection-zone (LPZ) of patients with macular degeneration (Masuda et al., 2021). Collectively, these fMRI data suggest that some form of cross-modal plasticity is possible in visual cortex.

Whilst transient disruption of visual cortex via TMS impairs Braille reading performance in blind individuals (Cohen et al., 1997), its detrimental impact appears to depend on the onset of blindness, with little impact on Braille reading performance of individuals whose blindness occurs after ∼14 years of age (Cohen et al., 1997, 1999). Thus, despite considerable fMRI evidence suggesting visual cortex is capable of cross-modal plasticity, whether or not such activity is causally related to cross-modal performance is less clear and may depend on plasticity of the brain that is only present early in life.

Prior fMRI work (Cheung et al., 2009), capitalized on the rare case of ‘patient S’, who despite being visually impaired, is capable of both reading print visually and Braille by touch. In S, tactile processing (e.g. Braille reading) selectivity recruited the foveal representation of visual cortex whereas visual processing (e.g. viewing letter strings) recruited more peripheral portions. There was no evidence of central-visual field loss in S, despite the loss of visual responses in the foveal representation. The fact that the foveal representation was recruited during Braille reading in S was interpreted as reflecting retinotopically specific cross-modal plasticity. Although it was argued that in S, such reorganisation was optimal - since only those parts of visual cortex that were not critical for S’s remaining low-vision were recruited during somatosensory processing - whether or not this somatosensory activity plays a causal role in S’s tactile processing ability is unclear.

Here, we tested this prediction directly in S by pairing both tactile and visual psychophysics with rTMS of the foveal representation of visual cortex (occipital pole [OP]), somatosensory cortex (S1) and an occipital lobe control region (OC). First, our fMRI experiment replicated prior work in S by demonstrating preferential recruitment of the foveal and peripheral representations of visual cortex during tactile and visual stimulation, respectively (Cheung et al., 2009). Second, we report that despite the pattern of fMRI data in S, transient disruption of OP via repetitive TMS (rTMS) does not alter tactile performance beyond that observed in normally sighted controls. The somatosensory-related activity within the foveal representation of visual cortex of S likely reflects unmasked latent connections with somatosensory cortex rather than reflecting causally relevant cross-modal reorganisation.

## Methods

### Participants

We report fMRI and behavioural measurements from 1 participant with ‘low-vision’ (Patient S; see **Case Description**) and three control participants with normal vision (C1-3; 2 male and 1 female; ages 21-35. Recruitment of low-vision patients who can still read visual print and are also expert Braille readers for basic research is difficult. Prior work, on which this study is based, published in *Current Biology*, focused on a case study of Patient S and compared fMRI response in S to those of 4 healthy controls. In the current case study of S, we have therefore adopted methods that enable us to perform measurements of single participants. All procedures adhered to protocols based upon the declaration of Helsinki ethical principles for research involving human participants. The ethics committees at the York Neuroimaging Centre and the Department of Psychology at the University of York approved these experiments. All participants provided written informed consent to participate in the experiment.

### MR Tactile and visual stimuli

Tactile stimuli in the form of Braille letters [(a, l, q or x)] were delivered via piezoelectric stimulator (max amplitude, 300ms). Eight presentations occurred during each 12s block. Visual stimuli consisted of 100% contrast reversing ring patterns (radial frequency 0.16 cycles per degree, reversal rate 6Hz). Ring stimuli extended to 15deg eccentricity. Each run consisted of 10 cycles of 12s on 12s off stimulation using an interleaved paradigm (Visual, rest, Tactile, rest).

### Scanning Procedure

All MRI data were acquired on a 3.0 Tesla GE Sigma HD Excite scanner. For structural data, two multi-average, whole-head T1-weighted anatomical volumes were acquired for each subject (repetition time = 7.8 ms, echo time = 3 ms, TI = 450 ms, field of view = 290 × 290 × 176, 256 × 256 × 176 matrix, flip angle = 20°, 1.13 × 1.13 × 1.0 mm3). For functional data, gradient recalled echo pulse sequences were used to measure T2* blood oxygen level–dependent data (repetition time = 2,000 ms, echo time = 30 ms, field of view = 192 cm, 64 × 64 matrix, 39 contiguous slices with 3-mm thickness). Images were read-out using an echo planar imaging (EPI) sequence. Magnetization was allowed to reach a steady state by discarding the first five volumes.

### fMRI Data Analysis and Visualisation

All anatomical and functional data were pre-processed and analysed using the Analysis of Functional NeuroImages (AFNI) software (Cox, 1996) (RRID: <u>SCR_005927</u>). All images were motion-corrected to the first volume of the first run (using the AFNI function *3dVolreg*). Following motion correction, images were detrended (*3dDetrend*) and spatially smoothed (*3dmerge*) with a 3mm full-width-half-maximum smoothing kernel. Signal amplitudes were then converted into percent signal change (*3dTstat*). To analyse the functional data, we employed a general linear model implemented in AFNI (*3dDeconvolve, 3dREMLfit*). The data at each time point were treated as the sum of all effects thought to be present at that time and the time-series was compared against a Generalized Least Squares (GSLQ) model fit with REML estimation of the temporal auto-correlation structure. Responses were modelled by convolving a standard gamma function with a 12s square wave for each stimulus block (Visual, Tactile). Estimated motion parameters were included as additional regressors of no-interest and fourth-order polynomials were included to account for slow drifts in the MR signal over time. To derive the response magnitude per condition, *t*-tests were performed between the condition-specific beta estimates and baseline. The corresponding statistical parametric maps were aligned to the T1 obtained within the same session by calculating an affine transformation (*3dAllineate*) between the motion-corrected EPIs and the anatomical image and applying the resulting transformation matrices to the T1. In each participant, the pre-processed functional data were projected onto surface reconstructions (*3dvol2surf*) of each individual participant’s hemispheres derived from the Freesurfer4 autorecon script (http://surfer.nmr.mgh.harvard.edu/) using the Surface Mapping with AFNI (SUMA) software (Saad & Reynolds, 2012).

### TMS target localisation

TMS target locations were defined in each participant individually. The occipital pole [OP] target was defined according to the T1-weighted anatomical scan. The occipital control [OC] target was defined as a fixed distance (∼1cm) dorsal and anterior of that participants’ OP target. Our team has employed a similar approach previously in order to define close proximity control locations relative to our primary target sites (Silson et al., 2013; Strong et al., 2017). The S1 target site was defined as the voxel showing the largest response to tactile stimulation within the appropriate portion of the somatosensory cortex.

### Psychophysical tasks and stimuli

Tactile and visual thresholds were established in each individual participant prior to TMS sessions. Note that due to S’s low-vision the size of the visual stimuli differed from controls. **Tactile threshold:** Braille letters (a, l, q or x) were delivered via piezoelectric stimulator. Each stimulus comprised all 6 pins, which were raised to a minimum pedestal level of 2250 (max available 4095) units. All pins were raised for 100ms, before a subset of these 6 pins were further raised to represent the Braille letter. Participants had to detect the target letter as the pins raised above the background ‘noise’. Using a 4AFC paradigm with a 1 up 2 down staircase, the maximum pin displacement was reduced while the noise pin amplitude was held constant to establish a 71% correct threshold for letter detection. **Visual threshold:** Maximum luminance visual letters (white, A, L, Q or X) were presented on a black background (15 degrees for S, 4 degrees for controls). Using a 4AFC paradigm with a 1 up 2 down staircase, the background luminance was increased while the letter luminance was held constant to establish a 71% correct threshold for letter detection.

### TMS Protocol

A train of four biphasic (equal relative amplitude) TMS pulses, separated by 50 ms (20 Hz) at 70% of the maximum stimulator output (2.6 T) were applied to the participants’ scalp using a figure-of-eight coil (50-mm external diameter of each ring) connected to a Magstim Rapid2 stimulator (Magstim). Participants were seated in a purpose-built chair with chin rest and forehead support. The coil was secured mechanically and placed directly above each cortical target (occipital pole [OP}, occipital control [OC], somatosensory cortex [S1]) with the handle oriented parallel with the floor. The position of the coil was monitored and tracked in real time allowing the displacement between the intended and actual site of rTMS delivery to be measured. Each participant underwent eight sessions (2 tasks × (3 TMS sites + 1 no TMS)). Each TMS session contained 35 trials (5 training). Stimuli (both Tactile and Visual) were presented according to each participants’ specific threshold. rTMS pulses were delivered concurrently with the presentation of the test stimulus. This temporal configuration was identical to that used in previous studies from our laboratory where induced functional deficits were found to be maximized when rTMS was delivered coincident with the stimulus onset.

### Resampling of rTMS data

Our study lacked the power to compare S’ behavioural performance to the average of the controls as is commonplace. Instead, we adopted a bootstrapping and resampling procedure to demonstrate that the impact of OP stimulation in S was not different from an expected distribution of controls. For each control and session, we randomly sampled 80% of the experimental data (24/30 trials) and calculated the percentage of correct responses. This procedure was then repeated 10,000 times before averaging these values across control participants. Distributions of the difference between conditions (e.g. S1 - OP tactile performance) were then created and compared with the same calculation in S.

### Subjective observations from S

We acquired subjective reports from S following rTMS sessions. Following S1 stimulation during tactile processing S reported that “all pins felt the same”, “Jaw reflex was a little distracting”. S did not report experiencing any tingling (i.e. tactile phosphenes) following S1 stimulation. Following OC stimulation during tactile processing, S reported that “it was a little easier than before (S1), but not great”. Following OP stimulation during tactile processing S reported that “That was easier than before (OC)”.

## Results

### Foveal recruitment during somatosensory processing

First, blood-oxygen-level-dependent (BOLD) fMRI was employed to localise tactile and visual responses in S and three normally sighted controls (C1-C3). **Figure 1**, shows the contrast of Visual *versus* Tactile overlaid onto surface reconstructions of both hemispheres for S (**Figure 1A)** and a representative control (C2, **Figure 1B**). In S, tactile responses are evident at the occipital pole in both hemispheres and throughout somatosensory cortex, whereas visual responses are restricted to more anterior portions of visual cortex that represent the periphery (Wandell et al., 2007). No such somatosensory related activity was observed in the visual cortex of C2. Indeed, visual and tactile responses were restricted to visual and somatosensory cortices, respectively - a pattern replicated in C1 and C3 **(Figure 1C)**.

**Figure 1.**
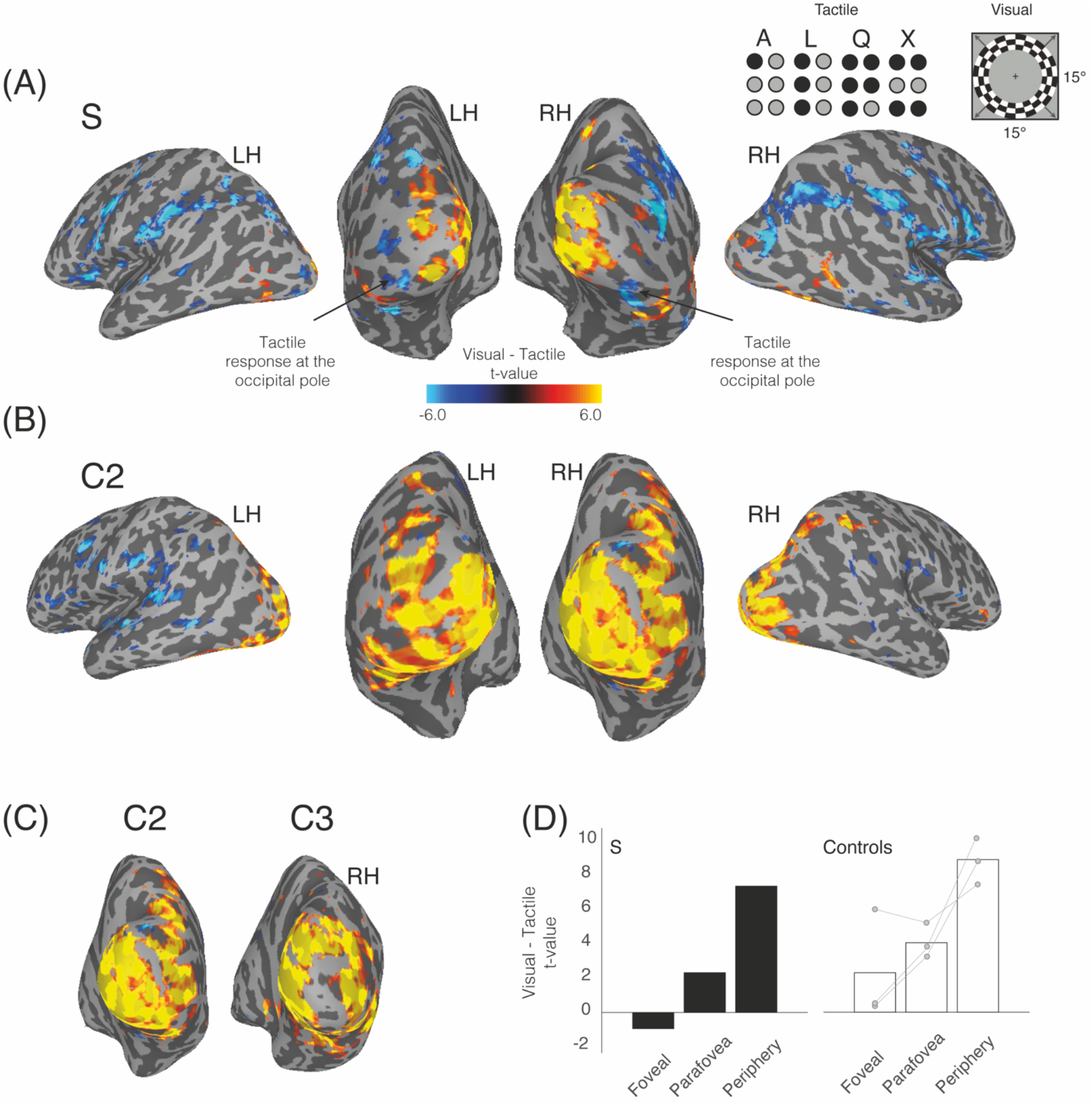
Tactile responses in foveal cortex of S. **(A)** Tactile and visual stimuli presented during fMRI are displayed inset. The contrast of Visual - Tactile is overlaid onto lateral and posterior partially inflated surface reconstructions of both hemispheres in S (LH=left hemisphere, RH=right hemisphere). Hot-colours represent visually evoked responses, cold-colours represent tactile evoked responses (*p*<0.0001, uncorrected). Tactile responses are evident at the occipital pole in both hemispheres. **(B)** Same as (A) but for a representative control (C2). No tactile responses are evident within visual cortex. **(C)** The contrast of Visual – Tactile is overlaid onto posterior views of the right hemisphere in the additional control participants (C1, C3). No tactile responses are evident at the occipital pole or within visual cortex. **(D)** Bars represent the mean response (t-value) within foveal, parafoveal and peripheral portions of early visual cortex in S (black bars) and the average of controls (white bars). Individual data points are plotted and linked for each control. Negative values represent larger responses during tactile processing, positive values represent larger responses during visual processing. In S, foveal cortex responds more to tactile over visual processing with the opposite pattern evident in parafoveal and peripheral portions. In controls, all portions show the anticipated larger responses during visual processing.

To confirm the replication of prior work (Cheung et al., 2009), three contiguous regions of interest (ROIs) were defined that divided primary visual cortex (V1) into foveal (< 4 deg eccentricity), parafoveal (> 4 < 8 deg) and peripheral (> 8deg) portions using eccentricity data from an independent group-average dataset derived from 29 healthy volunteers. **Figure 1D** shows the median response (given by the t-value for Visual *versus* Tactile) within each ROI for S and all three controls. In S, foveal responses are negative, reflecting tactile recruitment with both parafoveal and peripheral responses becoming increasingly positive (visual recruitment). In contrast, all three controls show positive responses, reflecting visual recruitment in all ROIs that increase in magnitude with increasing eccentricity.

### TMS target locations

**Figure 2A** shows the three TMS target locations in S, overlaid onto posterior and lateral partially inflated surface reconstructions of the right hemisphere, along with the contrast of Visual *versus* Tactile. The OP target site (black dot), which was defined according to the T1-weighted anatomical scan, can be seen to overlap the tactile responses (at the selected statistical threshold). The OC target (green dot) was defined as a fixed distance (∼1cm) dorsal and anterior of our OP target. Our team has employed a similar approach previously in order to define close proximity control locations relative to our primary target sites (Silson et al., 2013; Strong et al., 2017). The S1 target site (yellow dot) was defined as the voxel showing the largest response to tactile processing.

**Figure 2.**
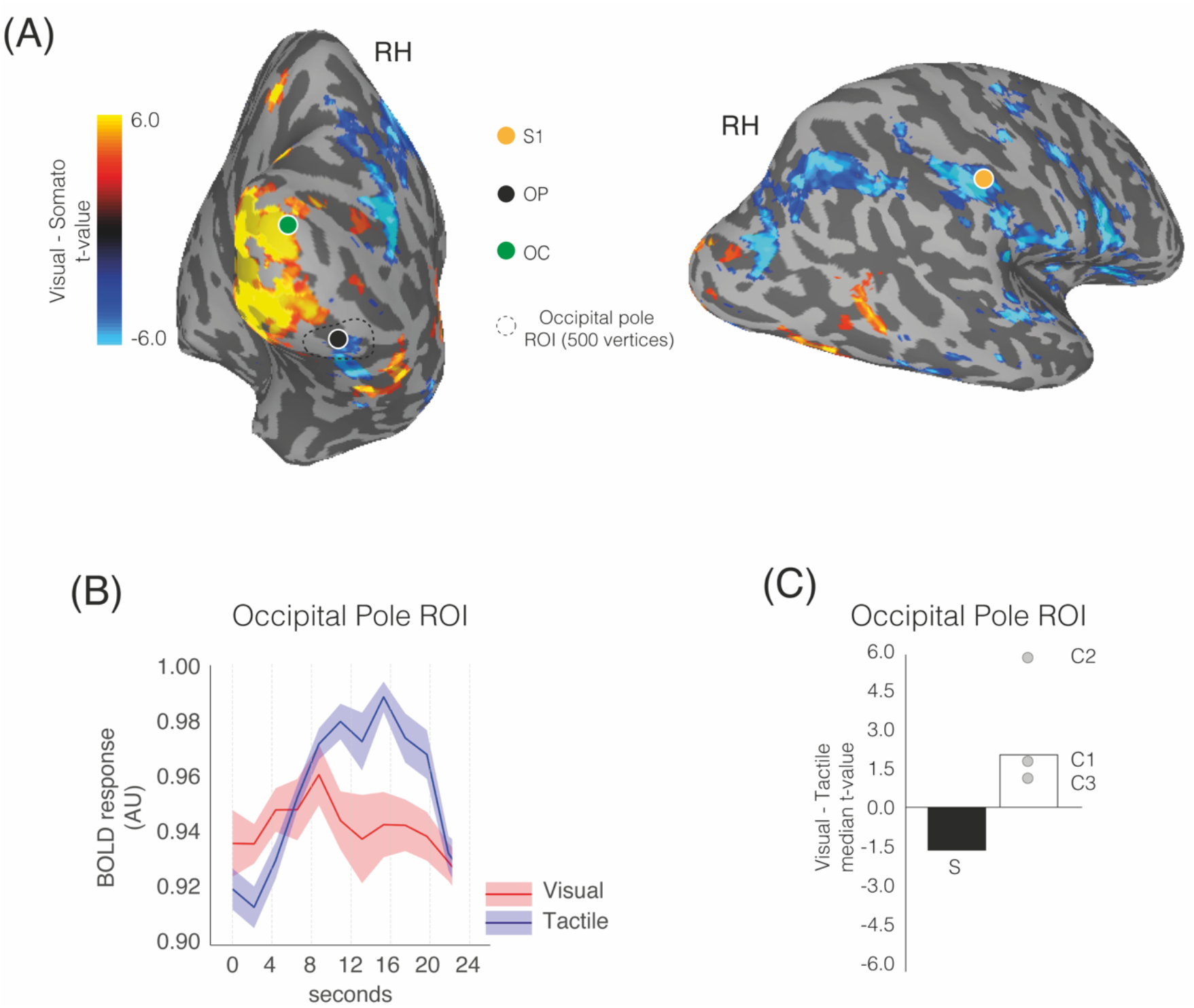
TMS target sites in S and fMRI responses from the occipital pole. **(A)** Posterior and lateral views of the right hemisphere of S are shown with the contrast of Visual - Tactile overlaid (*p*<0.0001, uncorrected). The occipital pole [OP] target site (black dot) can be seen to overlap tactile responses. The occipital control [OC] target site (green dot) is located dorsal and anterior of the OP. The somatosensory [S1] target site (yellow dot) can be seen to overlap tactile responses within the hand-representation of somatosensory cortex. The OP ROI encompassing the OP target site is shown by the black outline. **(B)** Lines represent the mean (plus s.e.m) response within the occipital pole ROI across all tactile (blue-line) and visual (red-line) fMRI blocks in S. The OP ROI selectively responds to tactile over visual processing. **(C)** Bars represent the mean response within the OP ROI in S (black bar) and the average of controls (white bars). Individual data points are plotted and labelled for each control. Whereas in S, the OP ROI shows a negative response reflecting selective recruitment during tactile processing, all three controls show the opposite pattern, reflecting the expected selective recruitment during visual processing.

To confirm that our anatomical OP target was within tactile responding cortex in S, we defined a region-of-interest (ROI) around the OP target site (500 vertices) and calculated the mean response from this ROI across all 10 tactile and visual fMRI blocks. **Figure 2B** shows the mean response (plus s.e.m) and highlights the preferential recruitment of this region during tactile processing in S. **Figure 2C** shows the median response (t-value) from this OP ROI for both S and all three controls. Whereas in S, a negative response is observed, reflecting tactile recruitment, the opposite pattern is observed in each control. Thus, the pattern of fMRI responses not only confirm prior work in S (Cheung et al., 2009), but also, highlight that responses in the foveal cortex of S are the opposite to those observed in normally sighted controls.

### rTMS of OP has no measurable impact on tactile processing performance in S

**Figure 3A** shows tactile performance in S and controls for all four conditions. These data reveal a strikingly similar pattern of performance across TMS conditions in S and controls - not predicted on the basis of the fMRI data. In S, tactile performance was maximally impaired (relative to noTMS baseline) following rTMS of S1 and to a lesser extent OC. Critically however, rTMS of OP had little to no detrimental impact on S’s tactile performance compared to the no TMS condition. On average controls showed a largely similar pattern with tactile performance maximally impaired following S1 stimulation but no clear detrimental impact of either OP or OC stimulation. Bootstrapping analyses indicate the impact of OP relative to S1 stimulation on tactile performance in S fell within the expected distribution of results in controls **(Figure 3B)**. Similarly, the impairment in tactile processing induced by TMS of S1 in S relative to noTMS baseline was of a similar magnitude to what could be expected from controls (**Figure 3C**).

**Figure 3.**
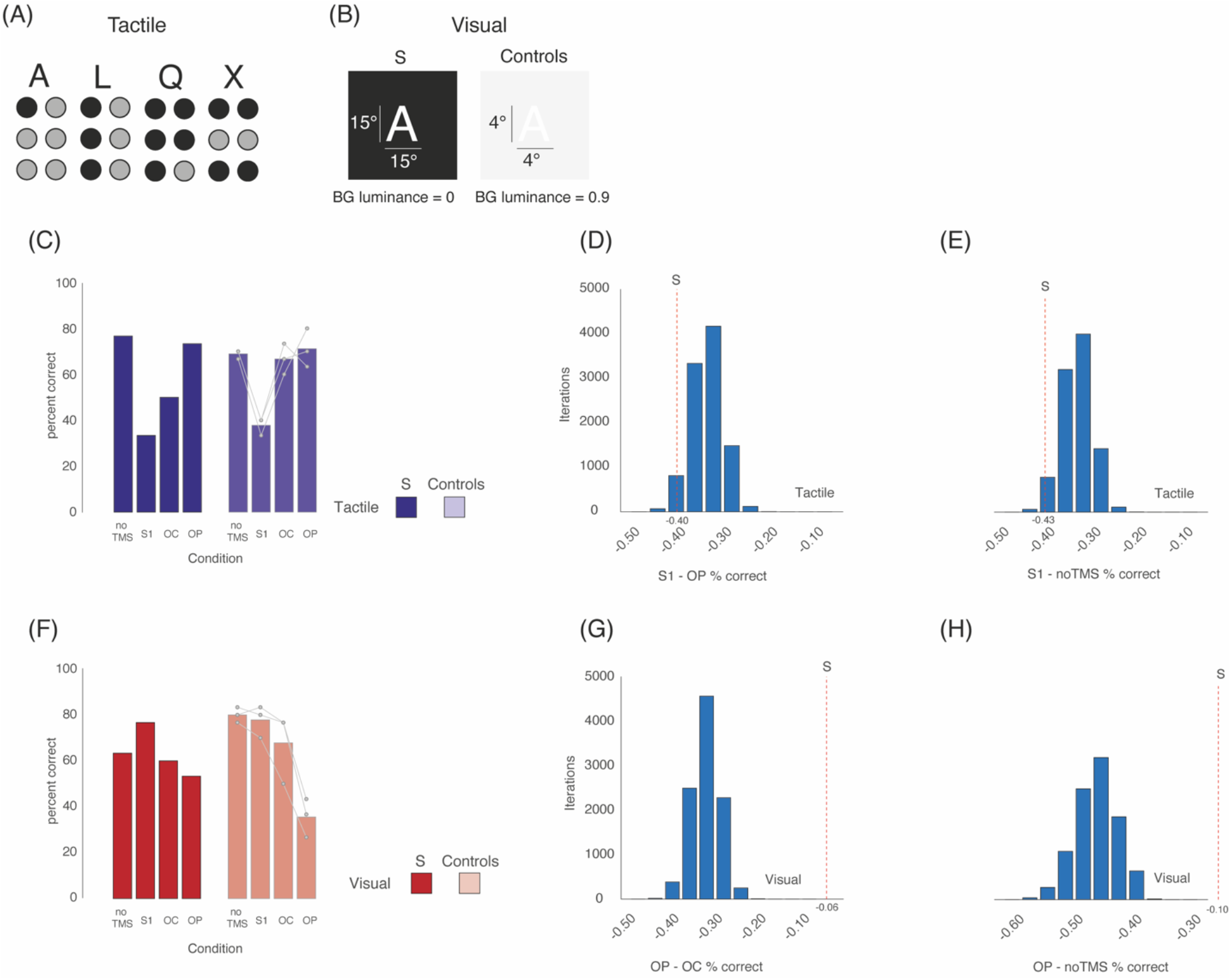
Impact of TMS on tactile and visual performances in S and controls. **(A)** Examples of the Braille letters presented during both psychophysics and rTMS sessions. **(B)** Schematic of visual stimuli presented during both psychophysics and rTMS sessions. During S’s rTMS sessions, letters were highly magnified (15 dva), white and presented on a black background (BG luminance = 0). In controls, letters were smaller (4 dva), and presented on a background luminance determined through psychophysical testing. Note, the figure indicates a BG luminance of 0.9 for illustrative purposed only (see Supplementary data for BG luminance thresholds in C1-C3). **(C)** Bars represent tactile performance (% correct) across conditions (noTMS, S1, OC & OP) in S (solid bars) and the average of controls (faded bars). Individual data points are plotted and linked for each control. The pattern of results is strikingly similar across S and controls. In both, performance is maximally disrupted (relative to noTMS baseline) following TMS of S1, as expected. In S, rTMS of OC caused a slight drop in performance, but crucially rTMS of OP had little to no impact on tactile performance in either S or controls. **(D)** Distribution represents the bootstrapped difference in performance between TMS of S1 - OP in controls (negative values represent a larger drop in performance following rTMS of S1). The red-dashed line indicates the same difference in S. Crucially, this difference falls not only within the distribution of expected differences from controls, but also towards the left-hand edge of the distribution (i.e. the maximum difference that could be expected in controls). This reflects the fact that the impact of OP stimulation in S on tactile performance is as small as could be reasonably anticipated in controls **(E)** Same as (D) but for S1 - noTMS baseline. Again, the result in S falls within that expected from controls. **(F)** Bars represent visual performance (% correct) across conditions (noTMS, S1, OC & OP) in S (solid bars) and the average of controls (faded bars). Individual data points are plotted and linked for each control. Unlike tactile performance, the pattern of results is more varied between S and controls. In both, performance is maximally disrupted (relative to noTMS baseline) following TMS of OP, but the magnitude of this disruption is larger for controls than for S. In S, rTMS of S1 caused an increase in performance, but had little to no impact in controls. **(G)** Distribution represents the bootstrapped difference in performance between TMS of OP - OC in controls (negative values represent a larger drop in performance following rTMS of OP). The red-dashed line indicates the same difference in S. The difference observed in S falls beyond that expected in controls. **(H)** Same as (G) but for OP - noTMS baseline. Again, the result in S falls outside that expected from controls.

### Impact of TMS on visual processing performance in S and controls

**Figure 3D** shows visual performance in S and controls for all four conditions. In S, visual performance was impaired slightly (relative to noTMS baseline) following rTMS of both OP and OC, but not S1 (which caused a slight increase in performance). In controls, performance was severely impaired following OP stimulation with much smaller decreases following stimulation of OC and S1, respectively - as was predicted for foveally presented small letter stimuli. Bootstrapping analyses indicate that the effect of OP stimulation on S’s visual performance is smaller than what could be expected compared to both stimulation of OC (**Figure 3E)** and the noTMS baseline (**Figure 3F**). The differential impact of OP stimulation on visual performance between S and controls likely reflects the fact that in S visual stimuli were required to be very large (∼15 deg) extending much further into the periphery, whereas the targeted OP region represents foveal visual field positions.

## Discussion

Our measurements suggest that the somatosensory related activity within the foveal representation of visual cortex of S plays little to no causal role in S’s tactile processing performance, and more likely reflects unmasking of latent connections between the somatosensory and visual cortices that are typically suppressed in normally sighted individuals (Masuda et al., 2021).

The pattern of TMS results in S during Braille reading were largely indistinguishable from those of the control participants, with stimulation of S1 inducing the largest detrimental impact to somatosensory processing. Critically, OP stimulation in S did not induce the reduction in tactile performance predicted on the basis of the fMRI experiments conducted here and in prior work (Cheung et al., 2009). That the foveal confluence of S preferentially responds to tactile over visual information was confirmed and yet the TMS data suggest that such activity is not causally related to tactile performance. In this regard, the pattern of TMS results in S are consistent with those of individuals with late-onset blindness (Cohen et al., 1999) This prior work demonstrated that TMS of occipital cortex induced tactile deficits in both congenitally blind (Cohen et al., 1997) and early-blind individuals but not those whose blindness occurred after 14 years of age (Cohen et al., 1999), suggesting a critical time-frame in which functionally relevant reorganisation of visual cortex occurs. Although S’s loss of visual function began at approximately six years of age, and thus within that timeframe, he nevertheless retains visual function. Indeed, S has a full visual field with no evidence of a central scotoma despite the very low-resolution ventral vision (Cheung et al., 2009). It is possible that this preserved peripheral visual function or his age when he lost vision has prevented the foveal representation in visual cortex taking on a causal role in tactile performance as is clearly the case in congenitally and early-onset blind individuals (Cohen et al., 1997, 1999; Sadato et al., 2002).

The finding that tactile responses in the foveal cortex of S play little to no causal role in S’s tactile performance offer the possibility that such cortical resources remain capable of high-resolution visual analysis even in the absence of such an input from the retinogeniculate pathway (Cheung et al., 2009). It is possible therefore that S’s foveal representation could revert back to processing high-resolution visual analysis if such retinogeniculate inputs could be restored (Fine et al., 2003) - although prior sight-restorations studies offer mixed encouragement for this possibility (Fine et al., 2003; Ostrovsky et al., 2006). Recent work in patients with macular degeneration (Masuda et al., 2021) highlight the presence of both somatosensory and auditory related activity within the LPZ during a one-back task, but not a passive condition. Such activity in the LPZ is considered to be mediated by task-related feedback signals, rather than feedforward visual input. The pattern of fMRI results in S could be interpreted in a similar manner, in that tactile responses within the foveal representation could reflect task-related feedback from S1 (although distinguishing feedforward from feedback signals definitively with fMRI is challenging due to the sluggishness of the fMRI response). Nevertheless, it is possible that the reduced retinal input to foveal representations in both S and patients with macular degeneration leads to an unmasking of pre-existing connections between visual and other sensory cortices that are suppressed during normal vision (Cheung et al., 2009; Masuda et al., 2021). Only one form of tactile perception (i.e. Braille discrimination) was measured here, and although the pattern of rTMS results is striking, it is nevertheless possible that some other form of tactile function (e.g. texture perception, tactile acuity) might benefit from the tactile recruitment of foveal visual cortex.

At first glance, it may appear surprising that TMS of OP during visual processing induced a much smaller decrement to performance in S than in controls. We believe however that this is accounted for by considering the size of the visual stimuli presented to S with respect to the foveal visual field representation of the targeted OP region. Placed in this context, it is not altogether surprising that rTMS of S’s OP induced a weaker deficit than TMS of OP in controls. It is likely that were it possible to stimulate peripheral parts of V1 in S, a similar drop in performance would be observed to that of OP stimulation in controls during visual perception. Additionally, we considered whether differences in the accuracy of rTMS delivery could provide an alternative explanation for the pattern of results reported here in S, and the critical finding that TMS of OP does not impact tactile processing, in particular. To rule out this possibility, we analysed coil-displacement data acquired during each rTMS trial - an index of stimulation error. We found no evidence for significant variation is displacement as a function of either task or site. Thus, the lack of a detrimental impact on tactile processing following rTMS of OP in S cannot be attributed to poor precision during rTMS delivery.

In summary, our study of S demonstrates that whilst foveal portions of visual cortex respond preferentially to tactile over visual stimulation, such activity does not causally influence tactile processing performance. Although prior work interpreted S’s responses in the foveal representation as reflecting an optimal redistribution of cortical resources (Cheung et al., 2009), our data suggests this pattern likely reflects the unmasking of latent connections between visual and somatosensory cortex that are normally supressed by the feedforward visual input provided to foveal cortex of normally sighted individuals (Masuda et al., 2021). We add weight to the view that cortical responses in individuals with visual deficits that differ from those obtained from controls are not always a signature of functional reorganisation.

## Acknowledgements

‘S’ is author G.E.L.

## Supplemental Data

**Table 1.** Tactile and visual thresholds for S and controls.

| Participant | Tactile threshold delta<br>(amplitude units) | Visual threshold<br>(Background luminance / size dva) |
| --- | --- | --- |
| **S** | 440 | 0.000 / 15 |
| **C1** | 762 | 0.988 / 4 |
| **C2** | 731 | 0.985 / 4 |
| **C3** | 411 | 0.986 / 4 |

